# Modulation of perception by visual, auditory, and audiovisual reward predicting cues

**DOI:** 10.1101/2023.01.29.526087

**Authors:** Jessica Emily Antono, Arezoo Pooresmaeili

## Abstract

Rewards influence information processing in the primary sensory areas specialized to process stimuli from a specific sensory modality. In real life situations, we receive sensory inputs not only from one single modality, but stimuli are often multisensory. It is however not known whether the reward-driven modulation of perception follows the same principles when reward is cued through a single or multiple sensory modalities. We previously showed that task-irrelevant reward cues modulate perception both intra- as well as cross-modally, likely through a putative enhancement in the integration of the stimulus parts into a coherent object. In this study, we explicitly test this possibility by assessing whether reward enhances the integration of unisensory components of a multisensory object in accordance with the *supra-additive* principle of multisensory integration. Towards this aim, we designed a simple detection task using reward predicting cues that were either unisensory (auditory or visual, both above the detection threshold) or multisensory (audiovisual). We conducted two experiments, behavioral (experiment 1) and simultaneous behavioral and neuroimaging testing (experiment 2). We expected that reward speeds up reaction times in response to all stimulus configurations, and that additionally the reward effects in multisensory cues fulfill the *supra-additive* principle of multisensory integration. We observed that reward decreased response times in both experiments with the strongest effect found for the multisensory stimuli in experiment 1. However, this behavioral effect did not fulfill the *supra-additive* principle. Neuroimaging results demonstrated sensory supra-additivity at the classical areas involved in multisensory integration such as the Superior Temporal areas (STS), while reward modulation was found in the midbrain and fronto-parietal areas, reflecting the typical areas that receive dopaminergic projections. However, reward did not enhance the *supra-additivity* in the STS compared to a no reward condition. Instead, we observed that some of the reward-related areas showed a *sub-additive* modulation by rewards and areas exhibiting a weaker *supra-additive* response to audiovisual stimuli, namely the fusiform gyrus, were modulated by rewards of audiovisual stimuli as measured by a conjunction analysis. Overall, our results indicate that reward does not enhance the multisensory integration through a *supra-additive* rule. These findings inspire a model where reward and sensory integration processes are regulated by two independent mechanisms, where sensory information is integrated at an early stage in a *supra-additive* manner, while reward modulates perception at a later stage *sub-additively*. Moreover, an associative area in the Fusiform gyrus exhibits a convergence of both reward and multisensory integration signals, indicating that it may be a *hub* to integrate different types of signals including rewards to disambiguate the information from different sensory modalities.

## Introduction

When we are enjoying our environment, for instance in a park, we experience a multitude of rich sensory inputs, such as birds chirping, people chatting while walking their dog, the wind blowing, and so on. Although the stimuli we encounter are coming from different senses (e.g. we see the bird on the tree and we hear the bird chirps), we perceive them as a unity. In other words, the brain integrates multiple sensory signals into a single coherent percept.

This phenomena is called multisensory integration and has been extensively studied in the past (Calvert, Hansen, Iversen, & Brammer, 2001; Soto-Faraco, Kingstone, Calvert, Spence, & Stein, 2004; M. T. Wallace, Meredith, & Stein, 1993; Mark T. Wallace & Stein, 1997), originating from the evidence in the neurophysiological study by Meredith & Stein, (1986), where the neural activity in the Superior Colliculus (SC) of anesthetized cats in response stimuli from either auditory, visual, or audiovisual was measured. When the cat received audiovisual stimuli, the neurons in the SC showed a response that exceeded the sum of the responses to the unimodal stimuli. At the neural level, multisensory integration is defined operationally as a statistically significant difference between the number of impulses evoked by a multisensory cue and those evoked by the most effective of the unisensory cues (Stein & Stanford, 2008). This exceeded response is the hallmark of multisensory integration and is argued to underlie the mechanism by which the brain suppresses the noise evoked by each stimulus alone, thereby disambiguating the percept of a multisensory object (Diederich, 1995; Diederich & Colonius, 2004; Stein & Stanford, 2008). Computationally, multisensory integration is a phenomenon whereby a response to multisensory stimuli exceeds the pooled responses to the unisensory cues, referred as a *supra-additive* response, whereas when the multisensory response shows equal or less response than the pooled response to unisensory cues, it is referred as an *additive* or *sub-additive* response (Colonius & Diederich, 2017; Stein, Stanford, Ramachandran, Jr, & Rowland, 2009). The *supra-additivity* rule has been observed in both neural (Stein, Meredith, & Wallace, 1993; M. T. Wallace et al., 1993) and behavioral responses to multisensory stimuli (Diederich & Colonius, 2004). Critically, in order for multisensory integration to occur, two (or more) sources of stimuli have to coincide in time and in space, i.e., they should occur simultaneously and be located at the same spatial location (Otto, Dassy, & Mamassian, 2013; Stein & Stanford, 2008). Moreover, the more ambiguous/noisy the stimuli are, the better the cues will be integrated (i.e. higher neural response for multisensory cues), a phenomena called the inverse effectiveness (Diederich & Colonius, 2004; Otto et al., 2013; Stein & Stanford, 2008).

Although multisensory integration effect is quite robust, its underlying mechanism has been disputed. Some argue that multisensory integration occurs automatically, as it can happen without the involvement of cognitive factors such as attention or awareness (Romei, Murray, Merabet, & Thut, 2007), while other researchers have provided evidence for an involvement of cognitive factor such as attention (Talsma & Woldorff, 2005) and awareness (Delong & Noppeney, 2021; Lunghi, Verde, & Alais, 2017). At a neurophysiological level, the automaticity of multisensory integration effects has been supported by showing that these effects arise at the level of subcortical or primary sensory areas, as the studies using anesthetized cats have shown (M. T. Wallace et al., 1993). However, multisensory integration response has also been observed in other areas such as in the intraparietal, superior temporal, and frontal cortex (Calvert et al., 2001; Lewis & Van Essen, 2000; Linden, Grunewald, & Andersen, 1999; Schroeder & Foxe, 2002; Stein & Stanford, 2008). These areas are known to process multisensory cues, as they respond to more than one sensory input but are also involved in higher cognitive functions. This indicates that there is an interplay between cortical and subcortical areas regulating multisensory integration that engages different sites, depending on the context. This raises a question as to whether multisensory integration is indeed an automatic process or does it depend on higher cognition?

Previous studies have investigated the role of attention, as a form of top-down/cognitive factor, in multisensory integration. An event related potential (ERP) study found a larger amplitude in the frontal positivity for the attended multisensory cues compared to the unattended ones (Talsma & Woldorff, 2005). Similarly, Senkowski, Talsma, Herrmann, & Woldorff (2005) extended this finding by using gamma band response as a measure and found early (< 90ms) attentional modulation of multisensory integration. These studies showed that attention plays a role in the multisensory integration at an early stage of cortical signal processing. Furthermore, Degerman and colleagues (2007) showed that the underlying neural basis of attentional effects on multisensory integration is similar to the unimodal cues, where it involves areas in the frontal, temporal, parietal, and occipital cortex. Additionally, attended multisensory features produced a stronger response in the superior temporal areas, compared to the attended single feature of a multisensory cue. Moreover, Ferrari and Noppeney (2021) extended this finding using a combination of Bayesian modelling in psychophysics and neuroimaging evidence and proposed that attention guides multisensory perception by two distinct mechanisms: the pre-stimulus attention enhances the precision of the attended sensory inputs, while post-stimulus attention binds features into a coherent percept depending on whether the features need to be integrated or not. The common findings across all these studies is that higher cognitive factors such as attention play a critical role in the multisensory processes, and that rather than being automatic, multisensory integration results from an interplay between higher cognition and early sensory processing.

However, previous studies are largely limited to the attentional processes. Both attention and reward are cognitive factors that shape our behavior and guide our decision (Pessoa & Engelmann, 2010). Unlike attention, the role of reward in multisensory processing has not been much explored. Hence, there is little known about how generalizable the interaction between the higher cognition, such as reward, and the multisensory processes are. Recent studies have investigated the interaction of higher cognition and multisensory processes using reward, questioning the automaticity of multisensory integration as earlier studies in attention have done. For instance, Bruns, Maiworm, and Röder (2014) found that reward expectancy alters the audiovisual spatial integration processes. In their study, they observed that higher magnitude of monetary reward reduced the cross-modal binding between audio and visual cues in the *ventriloquism* effect (i.e. rewarded cues exerted smaller ventriloquism effect), as reward made the auditory cues more spatially separable. Their findings highlight that reward influences that multisensory processes that are thought to be automatic, where possibly the interaction between reward and multisensory processing is enabled through a mediation via cognitive control mechanisms. Furthermore, Bean, Stein, and Rowland (2021) also examined this question by investigating the role of reward associations in multisensory integration. They showed that irrespective of the complexity of the cues, animals approached multisensory objects more reliably than the unisensory objects. However, when the value associated to one of the unisensory components of a multisensory object did not match the other, this gain for multisensory cues was lost. Their study hence shows that reward is an important factor for an organism to bind sensory cues to each other. Specifically, when there is a congruent association between the cues, it is more likely that integration will occur. Similarly, a study by Cheng, Saglam, André, and Pooresmaeili, (2020) observing the reward-driven modulation of saccadic trajectories in human found that the congruency of reward value across unisensory cues determines the combined salience of multimodal stimuli. Moreover, Hoofs, Grahek, Boehler, and Krebs, (2022) investigated the interaction between reward and attention to alter multisensory perception. In their study, they found that multisensory rewarded cues had a qualitatively different modulation than orienting (i.e. attention) processes, where multisensory reward processes were not simply expressed as the sum of the unisensory responses. Instead, in an orienting task, they showed that visual cues had a stronger attentional capture than auditory cues which occurred very early, indicating an automatic attentional process. However, when reward was expected, this attentional capture by visual cues was reduced in that the simultaneous presence of auditory cues helped performance by employing a more strategic orienting to the cued location.

Previous studies above have shown that reward plays a role in multisensory integration (Bruns et al., 2014), where either the prior association of reward (Bean et al., 2021; Cheng et al., 2020) or reward probabilities (Hoofs et al., 2022) is critical for the integration process. However, it is not clear whether other properties of reward, such as the magnitude of predicted rewards also plays a role in the integration process. More generally, it is not known how and at what stage of processing reward and multisensory integration interact with each other. As areas that are responsive to the magnitude of reward such as the ventral striatum, have also shown multisensory integrative response (Reig & Silberberg, 2014; Stevenson, Kim, & James, 2009), one possibility is that reward-related areas are also responsible for the integration of multisensory cues when they are predictive of rewards. Another possibility is that reward signals have to be projected to the sensory association areas (e.g. STS) or even the primary sensory areas in order to be affect the multisensory integration. In this context, we have previously shown that unimodal reward-associated cues involved both modality-independent reward- and attention- related areas as well as modality-dependent sensory association areas such as the STS. In this study, we ask whether when the reward is signaled from multiple sensory modalities, its effect would engage yet another distinct processing pathway compared to unimodal rewards or whether it would concurrently engage areas that are involved in the processing of unimodal rewards.

As attention has been more extensively investigated, there has been a systematic proposal of how attention and multisensory processes interact. Koelewijn, Bronkhorst, and Theeuwes (2010) proposed three different frameworks based on audiovisual attention studies. We adapted this framework to effects of reward, assuming that reward effects on multisensory mechanisms will follow a similar pattern as attentional effects. The *early integration* model (**Figure 1A**) proposes that multisensory integration would not occur without the reward information or in other words reward gates the multisensory integration. Here, we expect that reward modulates the responses of unimodal cues at a very early stage and since multisensory integration combines the already modulated responses to rewards, reward effects in multisensory stimuli are *supra-additive*. The *late integration* proposes that reward and multisensory integration are two independent mechanisms (**Figure 1B**). Here, we expect that sensory integration will occur first, producing a *supra-additive* response. Then, reward effect may enhance the response further. Note that in this framework, reward effect may not be *supra-additive*, but rather *additive* or *sub-additive*, as reward effect occurs at a later stage. The last model is the *parallel processing* scheme, suggesting that multisensory integration takes place at multiple stages (**Figure 1C**). Depending on the resources available, multisensory integration may occur at an early or late stage. Here, we expect that reward and multisensory integration share similar processing in the brain, where both of them may occur simultaneously in either sensory cortices or associative cortices. Similar to model A, model C also posits that both reward and multisensory processes occur at the same stage, hence leading to *supra-additive* reward effects.

**Figure 1.**
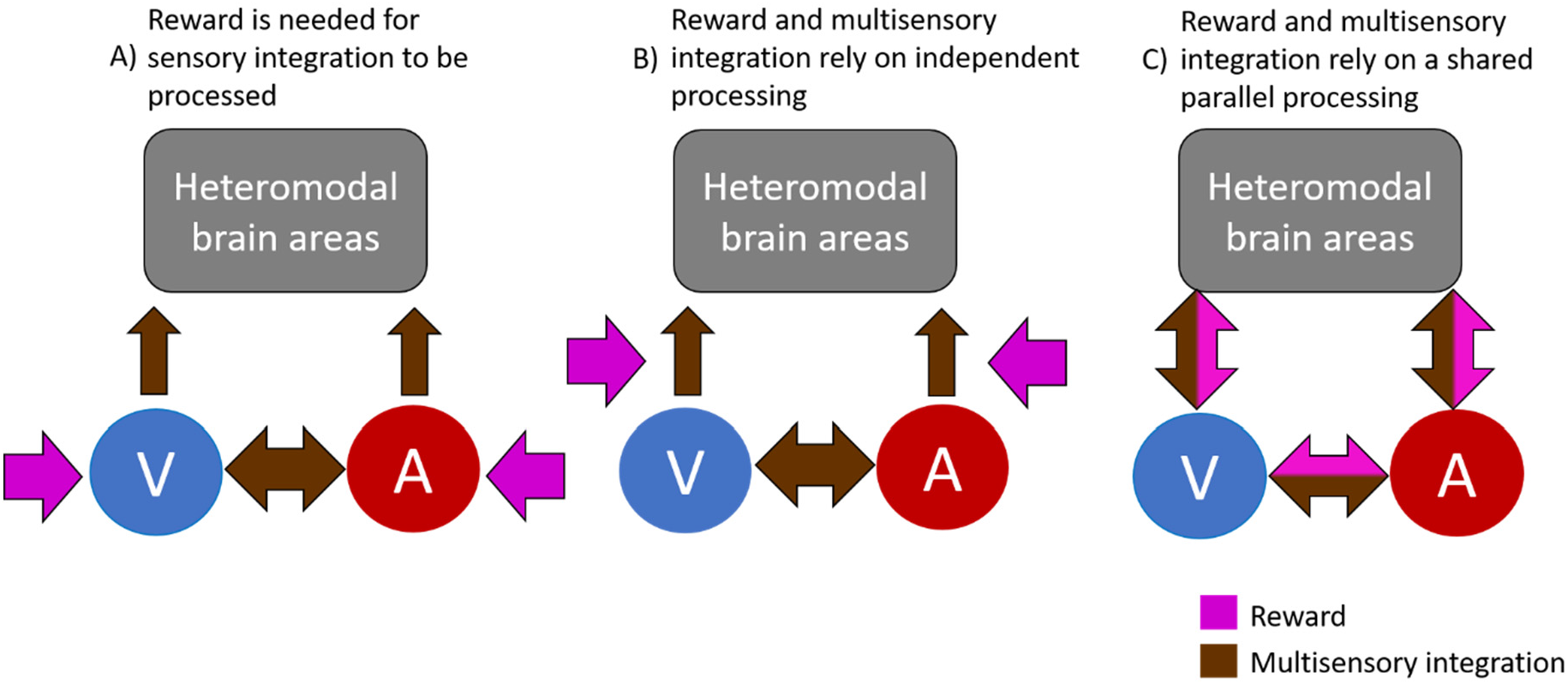
Schematic of the possible interactions between reward and audiovisual integration adapted from Koelewijn et al. (2010). In A), reward signal is needed in order for sensory cues to be integrated. Here, reward effect on multisensory integration is supra-additive. In contrast, B) proposes that reward and multisensory processes are two independent mechanisms, where multisensory integration occurs automatically, and then reward might modulate the effect further. This is expressed as sub-additive or additive reward effect in multisensory compared to unisensory stimuli. Alternatively, C) proposes that reward and multisensory integration are processed at multiple stages simultaneously. Since reward and multisensory integration occur at the same stage of processing, this model also predicts a supra-additive reward modulation for multisensory stimuli (but this supra-additivity could be observed at the level of heteromodal brain areas and not necessarily at the level of primary sensory areas).

In order to test the role of the reward in the multisensory integration, we designed a behavioral paradigm employing a detection task, where participants were asked to respond upon detection of unimodal or multimodal cues. In order to manipulate the reward magnitude, one feature of the cues, such as color or sound pitches, predicted reward where upon correct detection a reward was obtained, whereas another feature was paired with no reward. We recorded the behavioral responses -reaction times (*experiment 1 & 2*)-, and the behavioral and functional MRI (fMRI) responses simultaneously (*experiment 2*).

We expected that the reward effect will follow a different mechanism in multisensory compared to unisensory cues. Behaviorally, this distinction can be manifested as an interaction between the effect of reward and sensory modality on response times, where specifically the reward effect in multisensory cues significantly differs from the reward effect in the unisensory cues. Moreover, as our previous study (Chapter 3) showed that reward modulation is observed already at the level of early sensory areas, we expected that reward and multisensory processes also occur at the early stages of information processing. Hence, the underlying neural mechanism that might support this interaction is either the *early integration* or *parallel processing*. The common feature of these two mechanisms is that the reward signal is integrated at the same stage (or even earlier) as sensory cues are processed, hence producing a supra-additive reward-driven modulation of behavioral and neural responses.

Our results showed that although reward conferred a gain on performance across all unimodal and multimodal cues, its effect did not follow a supra-additive principle, as the enhancement gained from multimodal rewarded cues was similar to the one observed for unimodal rewards. Similarly, examination of the fMRI responses showed that reward effect in multimodal stimuli did not follow a supra-additive principle. However, we found evidence supporting *the late integration* model.

## Materials and Methods

### Participants

We collected data in two experiments, where in the first experiment only behavioural and eye tracking data were collected, and in the second experiment fMRI data was acquired. The target sample size for both experiments was set to by N = 20 before data collection started and was based on power calculation with a beta = 0.8 for the main effect of reward on auditory and visual modalities in 33 participants observed in a previous study (Chapter 3).

22 subjects participated in the *behavioral* experiment exclusively and 25 subjects participated in both *neuroimaging* including *behavioral* experiment. They were invited via an online recruiting system (http://www.probanden.eni-g.de/orsee/public/). All participants had normal or corrected-to-normal vision, had no history of neurophysiological or psychiatric disorders according to a self-report, and were naïve to the hypothesis of the project.

In the *behavioral* experiment, the total sample comprised 21 participants (14 male and 17 female, age 18 to 45 years; mean 25.54 years old ±5.36 years SD). 18 participants were right-handed and 3 participants were left-handed according to a self-report. 1 participant had to be removed from further analysis, as the participant’s accuracy in detecting the visual stimulus was below our inclusion criterion (accuracies < 70%). For two other participants, parts of the eye tracking data were not saved due to technical problem, nevertheless their data was analyzed to inspect the behavioral effects.

In the *neuroimaging* experiment, the total sample comprised 22 subjects (6 male and 16 female, age 18 to 45 years; mean 25.04 years old ±4.61 years SD). 20 participants were right-handed, while 2 participants were left-handed according to a self-report. Three participants were removed from our original sample (N = 25): one participant was outside of our age inclusion criteria (age > 45 years), one participant detected visual stimuli at ca. 45% accuracies, and for one participant data collection was not completed due to technical problems at the scanner.

Before the experiment started and after all procedures were explained, participants gave their oral and written consent. The study was approved by the local ethics committee of the “Universitätsmedizin Göttingen” (UMG), under the proposal number 15/7/15.

### Stimulus presentation and eye tracking apparatus

#### Behavioral experiment

Throughout the experiment, visual stimuli were displayed on a calibrated ViewPixx ASUS monitor subtending to 1080×1920 pixels, and a refresh rate of 120 Hz placed at a distance of 60 cm to the participants. For tracking the eye position an EyeLink 1000 Plus system with a desktop mount (SR Research) was used to track the right eye. The EyeLink camera was controlled by the corresponding EyeLink toolbox in MATLAB (Cornelissen, et al., 2002). Before each block, the eye tracking system was calibrated using a 9-point standard EyeLink calibration procedure.

The visual stimulus was a circle with a radius of 1.1° filled with a checkerboard pattern presented at 12° distance from the center fixation point. The colors of the checkerboard were orange and blue, RGB values are [255, 74, 44] and [0, 182, 155]. For auditory cues, two pure tone pitches (350 Hz or 1050 Hz) were presented at 70 dB. The tones were delivered through an over-ear headphone. In order to achieve the co-localization of auditory stimuli, we implemented a head-related transfer function based on a recorded database (Algazi et al., 2001) to localize the sound to be perceived as 12° distance relative to the center by taking into account dimensions of participant’s head measurement (i.e. width, height, depth, and circumference).

#### Neuroimaging experiment

Throughout the experiment, visual stimuli were displayed on an MR-compatible projection screen using a calibrated ProPixx projector (VPixx Technologies, Saint-Bruno, QC, Canada) at a resolution of 1920×1080 pixels, and a refresh rate of 120 Hz. The screen was placed at the end of the scanner bore at a distance of 88 cm from the participants’ eyes. The full display size on the screen was 43 cm x 24 cm, i.e. the visible range from the central fixation spot was +/- 13.6° horizontally and +/-7.7° vertically. For tracking the gaze position a ViewPoint eye-tracker system mounted on the mirror on top of the MR head coil was used (ViewPoint Eye Tracker, Arrington Research). Before the two scanning session, the eye-tracking system was calibrated using a 9-point standard ViewPoint calibration procedure.

The visual stimulus was a circle with a radius of 1.1° filled with a checkerboard pattern presented at 12° distance from the fixation point. In order to keep the same distance and ensure the visibility of the cues are captured within the screen display, we presented the fixation at 5.5° upwards relative from the center of the screen. The colors were isoluminant orange and blue, RGB values [255, 74, 44] and [0, 182, 155]. For auditory cues, two pure tone pitches (350 Hz or 1050 Hz) were presented at 90 dB. The tones were delivered through MR compatible earphones (Sensimetric S15, Sensimetrics Corporation, Gloucester, MA) with an eartip (Comply™ Foam Canal Tips) to maintain acoustic seal and reduce environmental noise.

### Procedure

The experiment consisted of a practice session (6 trials) for the simple detection task and two phases. In the first phase, one color and one pitch (e.g. color1, pitch1) were paired with reward. In the second phase, the reward association was switched, in that the color and the pitch that was predicting reward did not predict reward anymore in the second phase, but the other color and pitch (e.g. color2, pitch2) now predicted reward. This cue-reward association reversal was completed in order to counterbalance participants’ sensory bias due to the physical properties of the stimulus. The reward magnitude of the reward predicting cues was drawn from a Poisson distribution (mean = 25 Cents). Throughout the experiment participants had to respond upon detection of either a visual (90 trials), auditory (90 trials), or audiovisual (90 trials) stimulus. We also inserted empty trials (40 trials) to extend the inter-trial intervals randomly to reduce predictability of the onsets of stimulus. Following a response, participants received a feedback display: if participants had correctly detected the stimulus the feedback display showed their obtained reward (e.g. ‘20 Cent’ for rewarded and ‘00 Cent’ for not rewarded), and if they had missed the stimulus, no reward (‘00 Cent’) was shown. Importantly, we associated the same amount of reward across all cues, ensuring that the reward effect that we expect in the multisensory cues will not be due to a different magnitude of reward associated between unisensory and multisensory cues. The timing of events was identical across all phases. As soon as participants fixated (within 1° of the fixation point) a trial started. After a jittered fixation period of 3000-5000 ms (with a 1000 ms step), a target stimulus appeared (either a colored checkerboard for visual condition, a pure tone for auditory condition, or both checkerboard and the tone presented simultaneously for audiovisual condition). The target stimulus disappeared after 150 ms and participants had to press a button using their dominant hand to confirm detection within 2000 ms following the onset of the target. Finally, a feedback display was presented for 500 ms showing participants’ reward, as described above.

In order to determine whether participants learned the reward-cue association, they were asked to indicate which cue from each modality gave them more money. This question was completed in multiple parts following the first 30th, 60th, 90th trials and at the end of each phase (i.e. after 290 trials). Additionally, we also repeated the question in the questionnaire after the experiment was completed. All participants had learned reward associations based on their online results (in which at least one correct answer to the question was set as our criterion) and the questionnaire.

### fMRI data preprocessing for univariate analysis

Imaging data was processed using Statistical Parametric Mapping software (version SPM12: v7487; https://www.fil.ion.ucl.ac.uk/spm/). Preprocessing pipeline consisted of realignment of the slices to the mean image and unwarping the images according to the voxel displacement mapped image, slice time correction for multiband interleaved sequence, coregistration of the functional images to the structural scans, segmentation of brain structure according to the tissue probability maps, spatial normalization to the native space, and a spatial smoothing with a kernel size of 8 mm.

### Analysis of the behavioral data

The behavioral data obtained from all parts of the experiment was analyzed using custom-written scripts in MATLAB (version R2015a). We analyzed accuracies, reaction times (mean reaction time on a trial in which participants responded). We excluded the first 30 trials as participants needed time to learn the reward and cue association. Furthermore, we removed trials in which participants had reaction time shorter than 100 ms or larger than 2000 ms, or did not respond. This resulted in average 0.53% trials (±1.29 SD) for *experiment 1* and 0.36% trials (±1.41 SD) for *experiment 2* removed. For each response variable, we calculated the mean across all trials of each condition per subject during the pre- and post-reversal phases separately. Afterwards, the response variables (i.e. reaction times) was entered to a 2×3×2 repeated measures ANOVA (rm-ANOVA), with the reward (rewarded or not), sensory modalities of the cue (visual, auditory, or audiovisual), and phase (pre-reversal and post-reversal) as within-subjects factors. Then, we employed a Bonferroni corrected multiple comparison (using *multcompare* function in MATLAB) for the *post-hoc* test.

To quantify the performance gain of multimodal cues in relation to the unimodal gain, we measured the multisensory response enhancement (MRE). The MRE provides information on how much faster responses in the AV are relative to the fastest condition in the A or V, expressed as a percentage. When the values are positive, it indicates enhancement, while negative values demonstrate interference (Van der Stoep, Van der Stigchel, Van Engelen, Biesbroek, & Nijboer, 2019). Afterwards, we compared the MRE from reward predicting and not-reward predicting cues using a paired sample *t-test*.

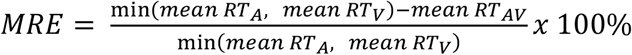

### Analysis of the fMRI data: Univariate analysis

For the univariate approach, we designed a General Linear Model (GLM) with regressors defined based on the reward magnitude factor (R-rewarded or NR-not rewarded) and sensory modality factor (V-visual, A-auditory, AV-audiovisual) which resulted in 6 regressors of interest as stick function. Moreover, we also modelled 8 nuisance regressors such as 6 movement parameters, events of no interest (e.g. instructions), and the phases of the task (pre- and post-reversal for each participant). For the univariate analysis, we entered preprocessed images to the General Linear Model (GLM). Following the estimation of each regressor in the GLM, we then defined a contrast of each 6 regressors of interest against baseline to be entered in the factorial design for group-level analysis.

In the factorial design, we defined a 2 by 3 repeated measures ANOVA with reward factor (rewarded or not rewarded) and sensory modality factor (audio, visual, and audiovisual) as within-subject factors and the participants were entered as the random factor. Then, T-contrasts were defined for the main effect of reward, main effect of sensory modality, and the interaction term. In the interaction term, we looked into the contrast of reward effect in each sensory modality against another configuration of sensory modality (e.g. Ar – Anr > Vr – Vnr; AVr – Avnr > Ar – Anr) in every possible combination (for a complete list of the contrasts, refer to **Table 1**). Moreover, we also explored reward effect in multisensory processes using an additional conjunction contrast between areas modulated by reward effect and *supra-additive* multisensory integration (main effect of reward ⋂ multisensory integration). Important to note, as the areas revealed by this conjunction responded to both *supra-additive* multisensory integration and reward effect, the neural response of these areas cannot be attributed to only one of the components (see Noppeney, 2012). Moreover, we also looked into the interaction contrast where reward effect audiovisual cues were stronger than the sum of reward effect in the unimodal cues (i.e. AVr - AVnr > ((Ar – Anr) + (Vr – Vnr))), as *supra-additive* reward effect, and also the vice versa reflecting the *sub-additive* reward effect. Overall, we thresholded the results with uncorrected *p* < 0.001 with extent threshold (*k*) of 10. The complete results of the whole-brain univariate analysis are shown in **Table 1**.

**Table 1.** Whole-brain activations of univariate results thresholded at uncorrected *p* < .001 and k = 10. Significance (*p*) showed at cluster-level. Activations marked in bold survived corrections for multiple comparisons at *p*FWE < 0.05.

| Cluster size | MNI coordinates (in mm) |  |  | T | p | Side | Region |
| --- | --- | --- | --- | --- | --- | --- | --- |
|  | x | y | z |  |  |  |  |
| Reward effect across sensory modalities ($AV_{rew}+A_{rew}+V_{rew} > AV_{not\ rew}+A_{not\ rew}+V_{not\ rew}$) | | | | | | | |
| 822 | 48 | 11 | 23 | 6.64 | < 0.001 | R | Inferior frontal operculum |
| 75 | 9 | 11 | -1 | 6.38 | 0.059 | R | Caudate |
| 631 | 54 | -34 | 53 | 6.20 | < 0.001 | R | Inferior parietal |
| 61 | -9 | 11 | -1 | 6.07 | 0.085 | L | Caudate |
| **295** | **-42** | **-37** | **41** | **5.99** | **0.001** | **L** | **Inferior parietal** |
| **292** | **42** | **-79** | **-7** | **5.69** | **0.001** | **R** | **Inferior occipital** |
| 43 | 6 | -1 | 29 | 5.33 | 0.142 | R | Cingulum |
| 81 | 9 | 26 | 44 | 5.10 | 0.051 | R | Frontal superior medial |
| **187** | **-48** | **-52** | **-13** | **5.03** | **0.006** | **L** | **Inferior temporal** |
| 78 | -30 | 17 | 2 | 5.01 | 0.055 | L | Insula |
| 143 | -42 | 5 | 23 | 4.83 | 0.013 | L | Frontal inferior operculum |
| 25 | -24 | -70 | -49 | 4.11 | 0.257 | L | Cerebelum |
| 17 | 9 | 38 | 14 | 3.92 | 0.350 | R | Anterior cingulum |
| 16 | 21 | 44 | -16 | 3.89 | 0.365 | R | Frontal superior orbital |
| 32 | 3 | -25 | 26 | 3.69 | 0.202 | R | Cingulum |
| 15 | -27 | -91 | -4 | 3.45 | 0.381 | L | Middle occipital |
| **Audiovisual reward effect ($AV_{rew} > AV_{not\ rew}$)** |  |  |  |  |  |  |  |
| 53 | 48 | -43 | 59 | 3.93 | 0.106 | R | Superior parietal |
| 43 | 33 | -91 | -1 | 3.79 | 0.142 | R | Middle occipital |
| 18 | 51 | 38 | 26 | 3.62 | 0.336 | R | Frontal inferior triangularis |
| 12 | 48 | 11 | 23 | 3.50 | 0.435 | R | Frontal inferior operculum |
| **Visual reward effect ($V_{rew} > V_{not\ rew}$)** |  |  |  |  |  |  |  |
| **70** | **-9** | **11** | **2** | **6.15** | **0.067** | **L** | **Caudate** |
| **595** | **48** | **38** | **17** | **5.41** | **< 0.001** | **R** | **Middle frontal** |
| **454** | **45** | **-34** | **44** | **5.12** | **< 0.001** | **R** | **Supramarginal** |
| 41 | 9 | 11 | 2 | 4.93 | 0.151 | R | Caudate |
| 105 | 33 | 20 | -1 | 4.52 | 0.029 | R | Insula |
| 97 | 9 | 26 | 44 | 4.46 | 0.035 | R | Frontal superior medial |
| 131 | 51 | -46 | -16 | 4.44 | 0.017 | R | Inferior temporal |
| 47 | -42 | -37 | 41 | 4.43 | 0.126 | L | Inferior parietal |
| 113 | -45 | -55 | -13 | 4.39 | 0.024 | L | Inferior temporal |
| 38 | -30 | 17 | 2 | 4.34 | 0.166 | L | Insula |
| 36 | -42 | 5 | 23 | 3.82 | 0.177 | L | Frontal inferior operculum |
| 30 | -24 | -64 | 44 | 3.74 | 0.216 | L | Superior parietal |
| 11 | 6 | -1 | 29 | 3.70 | 0.456 | R | Middle cingulum |
| **Auditory reward effect ($A_{\text{rew}} > A_{\text{not rew}}$)** |  |  |  |  |  |  |  |
| 39 | 9 | 11 | 2 | 4.55 | 0.161 | R | Caudate |
| 81 | 45 | 11 | 23 | 4.24 | 0.051 | R | Inferior frontal triangularis |
| 88 | 45 | 38 | 11 | 4.06 | 0.043 | R | Inferior frontal triangularis |
| 38 | 57 | -28 | 53 | 4.01 | 0.166 | R | Inferior parietal |
| 26 | 33 | 17 | -4 | 3.75 | 0.248 | R | Insula |
| 15 | -60 | -7 | 41 | 3.73 | 0.381 | L | Postcentral |
| 12 | 42 | -79 | -10 | 3.67 | 0.435 | R | Inferior occipital |
| 22 | -45 | -37 | 41 | 3.64 | 0.287 | L | Inferior parietal |
| 18 | -42 | 5 | 29 | 3.59 | 0.336 | L | Precentral |
| 11 | 3 | -31 | 29 | 3.38 | 0.456 | R | Middle cingulum |
| **Supra-additive multisensory integration ($AV > A + V$)** |  |  |  |  |  |  |  |
| **6561** | **63** | **-19** | **8** | **7.54** | **< 0.001** | **R** | **Superior temporal** |
| 162 | 21 | -85 | -43 | 4.93 | 0.009 | R | Cerebellum |
| 22 | 15 | 14 | 20 | 4.31 | 0.287 | R | Caudate |
| 79 | -18 | -88 | -40 | 4.24 | 0.054 | L | Cerebellum |
| 48 | -30 | -43 | -16 | 4.11 | 0.123 | L | Fusiform |
| 240 | -30 | 26 | 47 | 3.99 | 0.002 | L | Middle frontal |
| 33 | -15 | -67 | -19 | 3.93 | 0.195 | L | Cerebellum |
| 16 | 0 | -43 | 71 | 3.89 | 0.365 | L/R | Precuneus |
| 13 | -12 | 41 | -13 | 3.89 | 0.416 | L | Frontal medial orbital |
| 45 | -18 | -31 | 59 | 3.85 | 0.134 | L | Postcentral |
| 49 | 21 | 29 | 41 | 3.65 | 0.119 | R | Superior frontal |
| 41 | 24 | -31 | 53 | 3.62 | 0.151 | R | Postcentral |
| 30 | -42 | -73 | -46 | 3.61 | 0.216 | L | Cerebellum |
| 21 | 21 | -64 | -22 | 3.60 | 0.298 | R | Cerebellum |
| 21 | 48 | -58 | -40 | 3.59 | 0.298 | R | Cerebellum |
| **Sub-additive multisensory integration ($AV < A + V$)** |  |  |  |  |  |  |  |
| *No voxel survived* |  |  |  |  |  |  |  |
| **Interaction: auditory reward effect > visual reward effect ($A_{rew} - A_{not\ rew} > V_{rew} - V_{not\ rew}$)** |  |  |  |  |  |  |  |
| *No voxel survived* |  |  |  |  |  |  |  |
| **Interaction: visual reward effect > auditory reward effect ($V_{rew} - V_{not\ rew} > A_{rew} - A_{not\ rew}$)** |  |  |  |  |  |  |  |
| *No voxel survived* |  |  |  |  |  |  |  |
| **Interaction: audiovisual reward effect > visual reward effect ($AV_{rew} - AV_{not\ rew} > V_{rew} - V_{not\ rew}$)** |  |  |  |  |  |  |  |
| *No voxel survived* |  |  |  |  |  |  |  |
| **Interaction: audiovisual reward effect < visual reward effect ($V_{rew} - V_{not\ rew} > AV_{rew} - AV_{not\ rew}$)** |  |  |  |  |  |  |  |
| *No voxel survived* |  |  |  |  |  |  |  |
| **Interaction: audiovisual reward effect > auditory reward effect ($AV_{rew} - AV_{not\ rew} > A_{rew} - A_{not\ rew}$)** |  |  |  |  |  |  |  |
| *No voxel survived* |  |  |  |  |  |  |  |
| **Interaction: audiovisual reward effect < auditory reward effect ($A_{rew} - A_{not\ rew} > AV_{rew} - AV_{not\ rew}$)** |  |  |  |  |  |  |  |
| *No voxel survived* |  |  |  |  |  |  |  |
| **Interaction: supra-additive reward effect ($AV_{rew} - AV_{not\ rew} > (A_{rew} - A_{not\ rew}) + (V_{rew} - V_{not\ rew})$)** |  |  |  |  |  |  |  |
| *No voxel survived* |  |  |  |  |  |  |  |
| **Interaction: sub-additive reward effect ($AV_{rew} - AV_{not\ rew} < (A_{rew} - A_{not\ rew}) + (V_{rew} - V_{not\ rew})$)** |  |  |  |  |  |  |  |
| 25 | 9 | 11 | 2 | 4.08 | 0.257 | R | Caudate |
| 20 | -9 | 11 | 5 | 3.95 | 0.310 | L | Caudate |
| 15 | 51 | 5 | 26 | 3.53 | 0.381 | R | Inferior frontal operculum |
| 28 | 42 | 38 | 17 | 3.49 | 0.231 | R | Middle frontal |
| 12 | 9 | 38 | 14 | 3.48 | 0.435 | R | Anterior cingulum |
| 33 | 3 | 38 | 44 | 3.45 | 0.195 | R | Superior frontal medial |
| 13 | -48 | -43 | -16 | 3.44 | 0.416 | L | Inferior temporal |
| **Conjunction: main effect of reward ∩ supra-additive multisensory integration** |  |  |  |  |  |  |  |
| 56 | 42 | -58 | -19 | 3.72 | 0.098 | R | Fusiform |
| 14 | -54 | -1 | 38 | 3.53 | 0.397 | L | Precentral |

## Results

### Behavioral results: reward effect in multisensory cues was not stronger than unisensory cues

#### Experiment 1

We investigated reward effects when cued from unimodal (auditory and visual) or multimodal (audiovisual) stimuli. Participants detection accuracies were at a near perfect level (*audiovisual:* 99.61%, ±s.e.m 0.27; *auditory:* 99.66%, ±s.e.m 0.21; *visual:* 96.63%, ±s.e.m 0.91) and we focused our analysis on response times. A 2×3×2 repeated measures ANOVA on reaction times demonstrated a significant main effect of modality (F(2,40) = 71.38, *p* < 0.001, ηp^2^ = 0.78, see **Figure 3A**), where multimodal stimuli (mean = 367 ms, ±s.e.m = 15 ms) had the fastest response in comparison to both auditory (mean = 413 ms, ±s.e.m = 17 ms) and visual (mean = 457 ms, ±s.e.m = 13 ms) cues (all *ps* < 0.001). Moreover, reward predicting cues significantly sped up the response as observed in the reward effect across all sensory modalities (F(1,20) = 19.34, *p* < 0.001, ηp^2^ = 0.49, see **Figure 3B**) with the strongest reward effect observed in the post-reversal phase for multimodal cues (*p* = 0.03, Cohen’s d = 0.51), followed by a trend in the post-reversal phase for unimodal visual cues (*p* = 0.09, Cohen’s d = 0.38). These findings are partially aligned with our hypothesis, as the reward effect was strongest in the multisensory stimuli. However, the strong reward effect in multimodal stimuli did not reach a level to produce a statistically significant interaction effect between reward and modality factors (F(2,40) = 0.21, *p* = 0.76, ηp^2^ = 0.01), ruling out the hypothesis that reward effect would show different modulation when signalled from multisensory compared to unisensory cues. Moreover, as reward and cue associations were reversed, we observed a slower response time in the post- compared to pre-reversal phase (F(1,20) = 4.7, *p* = 0.04, ηp^2^ = 0.19), where the reaction time in the pre-reversal phase (mean = 406 ms, ±s.e.m = 14 ms) was faster than the post-reversal phase (mean = 418 ms, ±s.e.m = 16 ms), as expected as participants had to unlearn the previous associations and re-orient to the new reward and cue association. The rest of the effects were not significant (all *p*s > 0.1).

**Figure 2.**
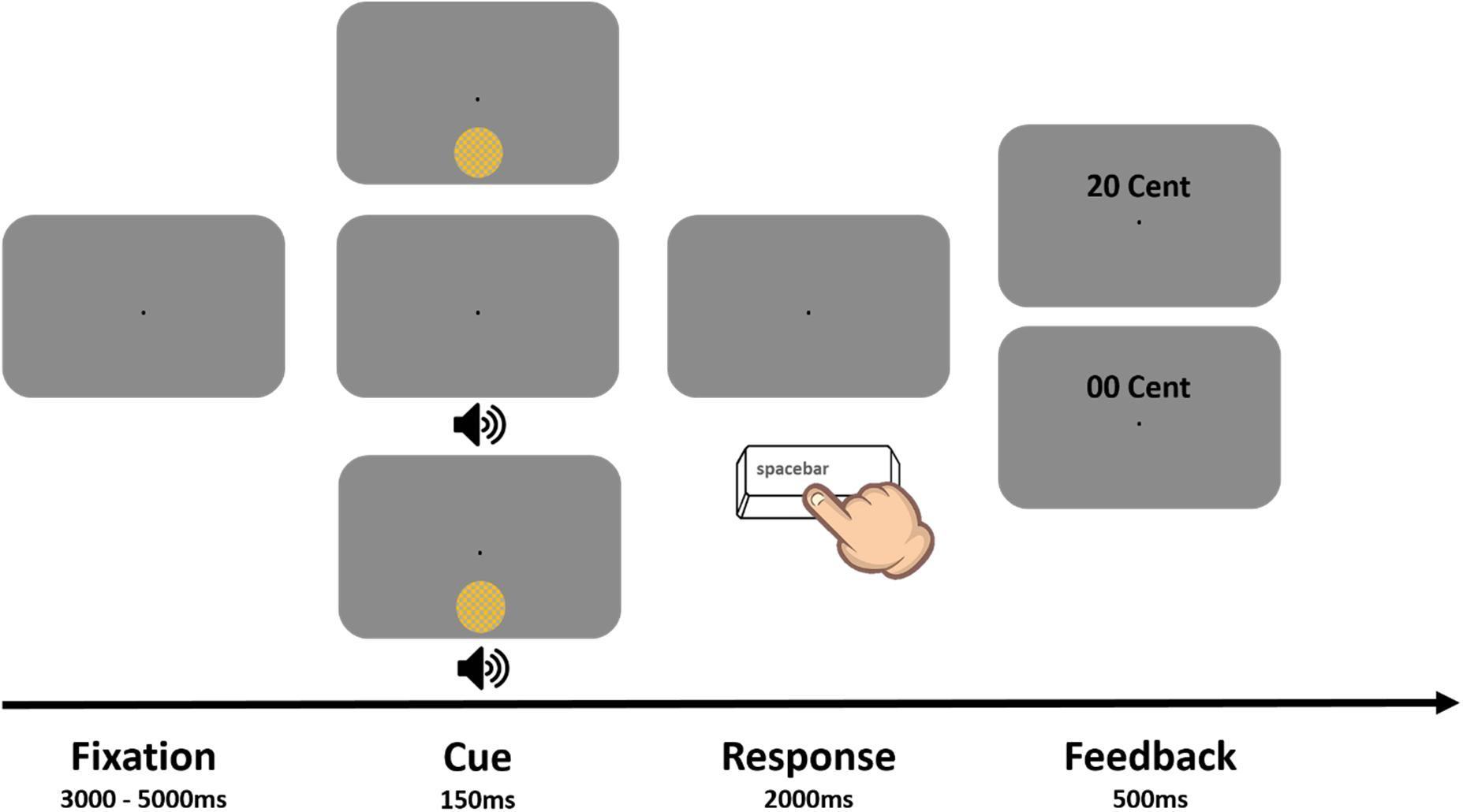
Behavioral paradigm of the simple detection task. Demonstration of the timeline of a trial, where upon fixation (3000, 4000, 5000 ms) either a visual, auditory, or audiovisual cue was briefly presented (150ms) in the periphery. Participants were asked to respond as fast as possible within 2000 ms response window by pressing a button. When detection was accurate, a feedback was displayed showing the reward receipt depending on the sensory features of the cue (e.g. blue color and high pitch tone predicted reward). If they missed a cue, ‘00 Cent’ was displayed. Reward associations of the cues were counterbalanced across participants. Furthermore, in order to remove any perceptual bias to cues that is unrelated to the reward assignments, we reversed the reward associations after halfway of the experiment.

**Figure 3.**
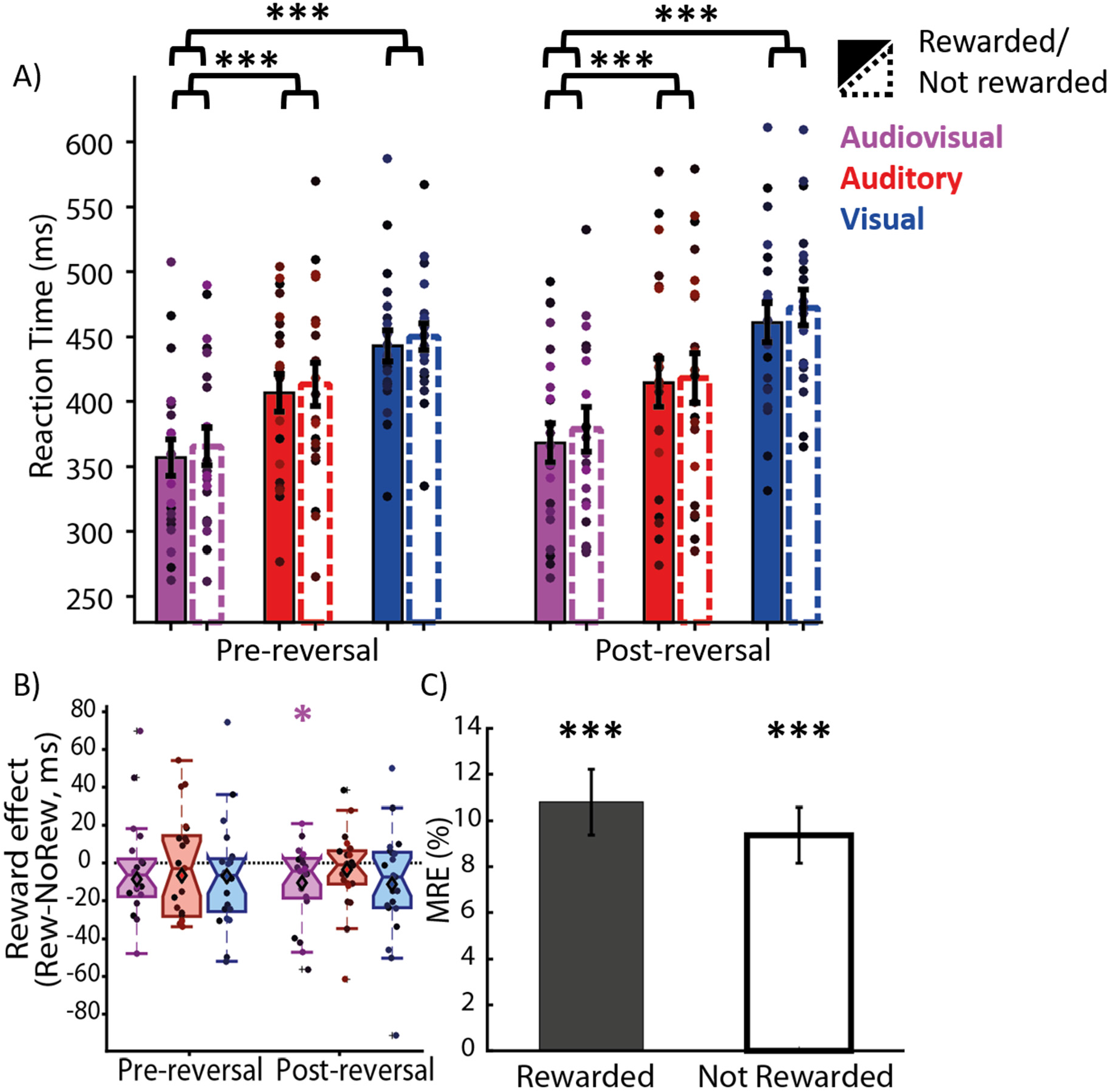
Experiment 1: Reward modulation of multisensory and unisensory cues. **A)** Bars depict the average response times for each condition at each phase. Colored circles correspond to the data of the individual subjects. Significance stars correspond to the effect of sensory modality. **B)** Reward effect in each sensory modality (auditory, visual, and audiovisual) across phases. **C)** Multisensory response enhancement (in %) between rewarded and not-rewarded conditions. * p < .05, *** p < .001.

However, to quantify how much gain a multimodal stimulus had over unimodal stimuli, we calculated the MRE for both rewarded and not rewarded stimuli (**Figure 3C**). We observed that multimodal cues enhanced the performance gain (rewarded: *p* < .001, Cohen’s d = 1.65; not-rewarded: *p* < .001, Cohen’s d = 1.68), pointing out that multisensory integration occurred. However, paired sampled *t*-test between rewarded and not-rewarded MREs did not reach significance (*p* = 0.28, Cohen’s d = 0.24), indicating that reward did not have a *supra-additive* effect.

#### Experiment 2

Similarly as experiment 1, we confirmed that participants’ detection accuracies were at a near perfect level (*audiovisual:* 99.89%, ±s.e.m 0.1; *auditory:* 99.59%, ±s.e.m 0.21; *visual:* 94%, ±s.e.m 1.26) and we focused our analysis on response times. We observed a main effect of modality (F(2,42) = 45.27, *p* < 0.001, ηp^2^ = 0.68, see **Figure 4A**), in which multimodal stimuli (mean = 368 ms, ±s.e.m = 21 ms) had the fastest responses in comparison to auditory (mean = 428 ms, ±s.e.m = 26 ms) or visual (mean = 441 ms, ±s.e.m = 18 ms) cues (all *p*s < 0.001). Also, rewarded cues made responses across all cues faster as observed in the main effect of reward (F(1,21) = 4.7, *p* = 0.04, ηp^2^ = 0.18, see **Figure 4B**). However, we did not observe any significant reward effect in the *post-hoc* tests examining individual modalities. There was also no interaction effect between reward and modality (F(2,42) = 2.36, *p* = 0.11, ηp^2^ = 0.1), indicating that reward in multisensory cues had a similar effect as in unisensory cues. The rest of the effects were not significant (all *p*s > 0.1).

**Figure 4.**
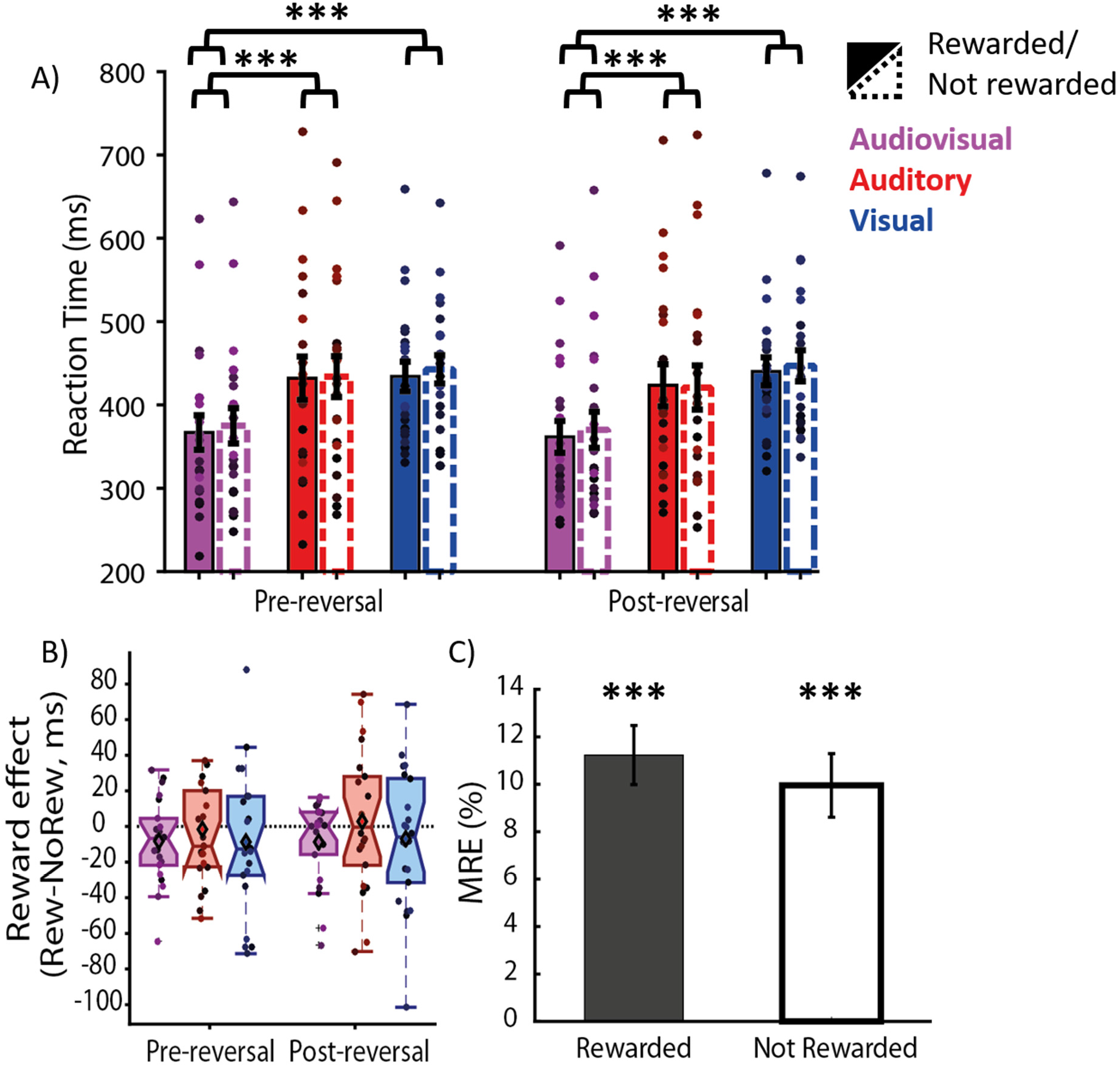
Experiment 2: Reward modulation of multisensory and unisensory cues. **A)** Bars depict the average response times for each condition at each phase. Colored circles correspond to the data of the individual subjects. Significance stars correspond to the effect of sensory modality. **B)** Reward effect in each sensory modality (auditory, visual, and audiovisual) across phases. **C)** Multisensory response enhancement (in %) between rewarded and not-rewarded conditions. * p < .05, *** p < .001.

Then, we investigated further the MRE, where here we also observed that even though multimodal cues enhanced the performance gain (rewarded: *p* < .001, Cohen’s d = 1.92; not-rewarded: *p* < .001, Cohen’s d = 1.58, see **Figure 4C**), reward did not make any significant difference (*p* = 0.18, paired t-test, Cohen’s d = 0.29). In general, although the strength of reward effect in experiment 2 was weaker than experiment 1, our behavioural results inside the scanner replicated the observations outside the scanner in terms of the reward effect and the fastest response observed in the multisensory cues. Collectively, the results from both experiments showed that reward effect in multisensory cues are similar to unisensory cues.

### fMRI results: whole-brain

Examining a contrast capturing the main effect of reward across sensory modalities, we observed reward modulations in the areas typically involved in the reward processing such as the striatum and frontal areas (**Figure 5A**). Moreover, we found the same areas for unimodal auditory and visual reward effects as shown in **Table 1**. We also tested the contrast of reward effect in each unimodal condition against the other (i.e. Ar – Anr > Vr – Vnr and vice versa) to ensure that the response profile in the unimodal conditions were similar. The reward and unimodal condition interaction contrasts did not reveal any modulation surviving the threshold, indicating that the reward effect is largely similar across sensory modalities. Furthermore, we also tested the interaction contrast of whether the reward effect in the multisensory cues was larger than the reward effect in each of the unisensory cues (i.e. AVr – AVnr > Vr – Vnr and AVr – AVnr > Ar – Anr, and vice versa for each contrast). In these interaction contrasts, we did not see any activation (**Table 1**), indicating that the reward effect was not larger in multisensory compared to the unisensory cues. As reward modulation might be expressed as an inhibition, we also looked into the reverse contrast across all conditions (Anr + Vnr + AVnr > Ar + Vr + AVr), where medial prefrontal areas, precuneus, and the temporo-parietal areas were recruited (see **Table S1**). The suppression of these areas by reward might indicate activities of some brain regions needed to be suppressed to enable reward-seeking behavior, as a previous study in mice observed that the modulation of the medial prefrontal cortex inhibited reward-seeking behavior (Ferenczi et al., 2016).

**Figure 5.**
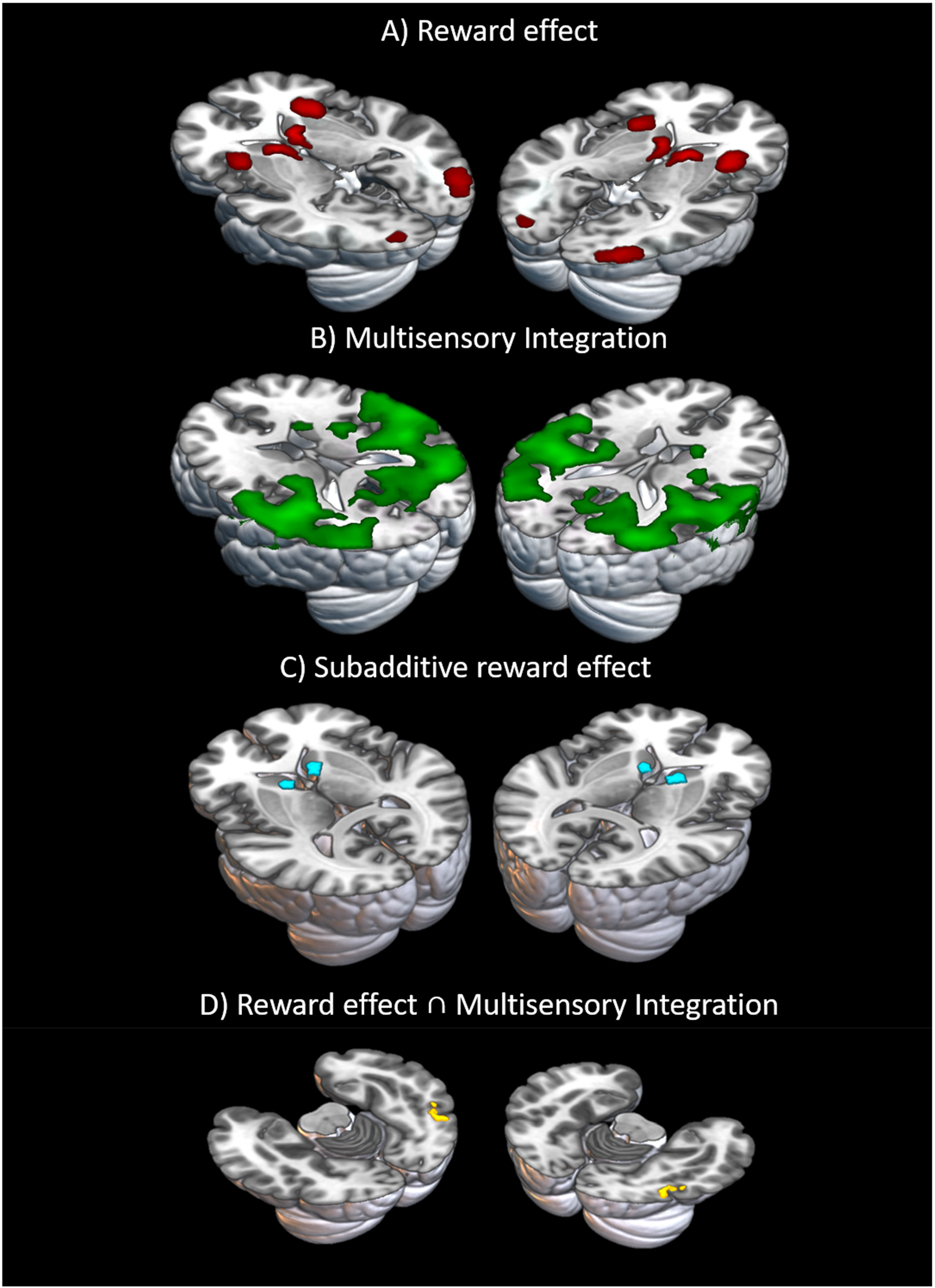
Experiment 2: whole-brain results. **A)** main effect of reward across sensory cues (in red), **B)** areas showing supra-additive responses to multisensory stimuli irrespective of the reward (in green), **C)** sub-additive reward effect, where the sum of reward effects in auditory and visual cues was higher than the reward effect in multisensory cues (in blue), **D)** and the results of a conjunction contrast between reward effect and sensory supra-additive responses to multisensory stimuli (in yellow). Note that we found no activities in the contrast capturing the supra-additive reward effects. All images were displayed at uncorrected p < .001 with extent threshold 10.

### fMRI results: multisensory integration effect, supra-additivity of responses to audio-visual stimuli and its modulation by reward

We next examined the areas that show supra-additive responses to multisensory stimuli (**Figure 5B**). To do this we tested the multisensory integration contrast (AV > A + V, across reward conditions). Here, we observed the strongest activations in the Superior temporal areas, as has been reported in previous studies (Beauchamp, 2005; Calvert et al., 2001). Other areas such as the fusiform gyrus and the precuneus were also showed supra-additive integration characteristics (**Table 1**). The observation in precuneus is in line with previous findings indicating that this area is involved in the multisensory processing (Renier et al., 2009) and fusiform areas have been also shown to be involved in binding of face and voice stimuli (De Gelder & Bertelson, 2003). Interestingly, the right caudate, an area strongly linked to the reward processing (Schultz, 2000), was also involved in integrating the multisensory cues showing a supra-additive effect, as has been also observed by previous studies (Nagy, Eördegh, Paróczy, Márkus, & Benedek, 2006; Reig & Silberberg, 2014; Stevenson et al., 2009). This might be an indication that reward processing might occur at the same stage as multisensory integration following a supra-additive principle. However, the sensory supra-additivity in Caudate was observed at locations that were distinct from regions that exhibited reward modulation (see **Figure S1**). Furthermore, to examine whether there are also areas responsible for a sub-additive integration, we looked into the sub-additive multisensory integration contrast (AV < A + V). However, we found no activation, indicating that the processing of the compound of the auditory and visual signals in the multisensory cues produces an effect that cannot be explained by the linear summation of the unisensory cues.

### fMRI results: examining the supra-additivity of reward effects in multisensory cues

The sensory integration contrast above showed that the multisensory cues had a different response compared to the unisensory cues. We hypothesized that the reward effects in multisensory cues would be different from the reward effects in unisensory cues. Therefore, to test this hypothesis, we looked into the *supra-additive* reward effect contrast (AVr – Avnr > ((Ar – Anr) + (Vr – Vnr)), **Table 1**). However, this contrast did not reveal any activation. In fact, areas with the maximum supra-additive response (i.e., STS: peak at xyz=[63 -19 8]) were not modulated by reward, as we looked specifically into the reward effect in the superior temporal areas (*p* = 0.47, Cohen’s d = 0.15, **Figure S2**). This finding could indicate that reward effect might enhance multisensory integration in a *sub-additive* manner, as has been observed before (Werner & Noppeney, 2010, 2011). Indeed, in the *sub-additive* contrast of the reward effect (AVr – Avnr < ((Ar – Anr) + (Vr – Vnr)), bilateral Caudates were modulated (right: [9 11 2]; left: [-9 11 5], **figure 5C**). This *sub-additivity* of reward effect might indicate that reward effect does not depend on the configuration of the stimulus. However, note that both our unimodal and multimodal cues were associated with the same magnitude of reward. In that case, contrasting the sum of reward effect in unimodal cues against reward effect in multimodal cues might reflect the processing of reward magnitude and not necessarily a specific interaction between the reward coding and multisensory integration.

Our evidence so far has been hinting towards independent mechanisms for the processing of reward and multisensory information, where sensory cues were integrated in a *supra-additive* manner in the sensory association areas, while reward information was processed as an additive factor (i.e. similar across uni- and multisensory cues) in the striatum. However, we asked further how the two processes converge; whether there is a region that acts as a hub to undertake both processes? In order to investigate this, we employed a conjunction analysis between the reward effect and multisensory integration contrast (i.e. main effect of reward ⋂ AV>A+V). The conjunction revealed activations in the Fusiform gyrus [42 -58 -19] and the premotor cortex [-54 -1 38] (see **Figure 5D** and **Table 1**). Furthermore, bilateral activations were observed in the Fusiform gyrus at a more lenient threshold (at xyz = [42 -58 - 19] as well as xyz = [-39 -61 -7], at *p* uncorrected < 0.005, k = 20). This result hence indicates that the Fusiform areas were modulated by both reward and sensory integration, indicating they might act as a hub where both reward and multisensory processing converge.

## Discussion

Our study aimed to investigate the effect of reward on the multisensory integration. In comparison to the unisensory reward effects, reward-driven effect on multisensory stimuli have been less explored, and thus it is unclear whether they are following similar principles or are regulated by distinct mechanisms. Previous studies have shown that multisensory cues elicit a distinct response compared to unisensory cues, adhering to a supra-additive principle. Therefore, we expected that if rewards influence the multisensory processes at an early stage in which sensory integration occurs (i.e., models A and C in **Figure 1**), their effect on the multisensory stimuli should also adhere to a supra-additive principle (i.e., be larger than the pooled reward-driven modulation of unisensory stimuli). Our behavioral results showed that although the sensory processing of the multisensory cues was distinct from the unisensory cues (i.e., supra-additivity in the speed of responses), the reward modulation did not show any distinction across the sensory modalities (i.e., a lack of an interaction between reward and sensory modality and supra-additivity). Moreover, our neuroimaging results showed similar effects. Although we found strong reward-driven modulations and supra-additivity in the responses to the multisensory stimuli, no interaction was found between the two. In fact, areas with the strongest supra-additivity for sensory integration, such as the STS, were not modulated by reward. Instead, we found two lines of evidence supporting a late integration model (panel B in **Figure 1**). Firstly, the responses of some of the reward-related areas such as the Caudate to multisensory rewards was smaller than the sum of their responses to unisensory rewards (sub-additivity), indicating that they are primarily sensitive to the reward magnitude and did not differentiate multi- and uni-sensory stimuli. Secondly, using a conjunction contrast, we found an area in the Fusiform gyrus showing both sensory supra-additivity and reward modulation, albeit no supra-additivity of reward effects. Therefore, this area might act as a convergence point between the reward and multisensory processing and contribute to the late integration of the two sources of information.

In the behavioral results, we observed that multisensory integration did occur, as indexed by the Multisensory Response Enhancement (MRE). However, we found no interaction effect between the sensory modalities and reward, indicating that multisensory integration and reward might be regulated independently or at different stages. This result did not resonate with our hypothesis that reward would be integrated as *early* as the sensory cues are. Furthermore, our results are in contrast with the observations from Bean and colleagues (2021), as in their study they observed a robust reward-driven enhancement of multisensory integration as measured by the approach behavior of cats. There are several differences in the paradigm used by the latter study and ours that might lead to this difference. First, the strength of the stimuli employed in the two studies was different. In our study, we employed supra-threshold cues presented at a single peripheral location, as we expected that irrespective of the strength of the stimuli, reward will enhance the multisensory integration. In contrast, the paradigm used in Bean and colleagues (2021), used low intensity unisensory cues presented at several randomized peripheral locations. Since according to the *inverse effectiveness* rule, multisensory integration is stronger for weaker unisensory stimuli, it is possible that reward only affects multisensory integration when unisensory stimuli are weak. In such settings, increasing the gain of the multisensory integration by rewards can have crucial behavioral advantages. Second, in our study, we did not manipulate the congruency of the reward in each of the unisensory stimuli, as both components of the bimodal cues predicted the same reward (i.e. either both visual and auditory cues were rewarded or unrewarded). In contrast, the studies from Bean and colleagues (2021) and Cheng and colleagues (2020) highlight the importance of the manipulation of the congruency between rewards of the unisensory cues. In their studies, reward congruency helped the brain to discern which unisensory stimuli had congruent rewards and hence were more likely to belong to the same object, similar to how spatial and temporal overlap promote the multisensory integration. In fact, as our paradigm did not manipulate the congruency of rewards in unisensory stimuli, there might have been no necessity for the organism to sort out whether the two unisensory cues in an audiovisual stimulus derive from the same source or not. However, we also note that using the reward congruency allows an additional contribution of semantic factors, i.e. the numeric or categorical value of rewards, to the observed effects. As we were primarily interested in the sensory and physical factors that drive the multisensory integration, we decided not to vary the reward congruency. However, future studies will be needed to determine the extent to which this factor plays a role in the reward-driven changes of multisensory integration.

Examination of the BOLD activities revealed a similar pattern to our behavioral results. Firstly, there were areas recruited in the multisensory integration process exhibiting a supra-additive response to the audiovisual stimuli, especially in the Superior Temporal areas (STS). STS has been consistently reported to converge and integrate signals from multiple sensory modalities (Beauchamp, 2005; Calvert et al., 2001; Degerman et al., 2007; Ferrari & Noppeney, 2021b). Secondly, we observed that reward enhanced the BOLD responses in the classical reward-related areas such as the caudate and the frontal areas. However, our index of supra-additivity of the reward effect did not show any activations. Together, these findings rule out our hypothesis that reward affects the multisensory integration in an early stage and through a supra-additive change in neuronal responses.

Further examination of the BOLD responses revealed several interesting findings. As we compared the reward effect of each sensory modality configuration using interaction contrasts, there was no difference in the reward effect depending on the sensory modality configuration. Indeed, the same coordinates in caudates showing significant activations for the main effect of reward, also revealed a *sub-additive* reward modulation for audiovisual stimuli, indicating that the reward modulation in this area is primarily sensitive to the magnitude of the reward (which was the same for all conditions) and does not differentiate multisensory cues from the unisensory stimuli. Together, these findings suggest that in the setting we employed the brain’s reward network is largely invariant to the sensory modality. Another interesting observation was that in the distinct regions of the caudates, an area primarily known for its role in reward processing, there were regions that also played a role in sensory integration (see **Figure S1**). Specifically, the right caudate showed a *supra-additive* response to multisensory stimuli compared to unisensory stimuli (across reward conditions). This is in line with previous studies, showing the existence of both unisensory and multisensory neurons found in the Caudate (Nagy et al., 2006; Nagy, Paróczy, Norita, & Benedek, 2005). Moreover, the Caudate has also been reported to be involved in the integration of sensory information in mice (Reig & Silberberg, 2014) and also humans (Stevenson et al., 2009). Hence, the Caudate has been known to respond to multisensory cues, as one of its critical function is to coordinate movement (Nagy et al., 2006). In our study, the peak of the Caudate that were modulated by reward effect and *sub-additive* reward effect in multisensory stimuli are similar, but the peak that was modulated by *supra-additive* sensory integration was distinct. Therefore, we conclude that the areas that receive sensory information in the Caudate do not overlap with the areas that respond to rewards.

Up to this point, our evidence pointed that reward and sensory integration are two independent mechanisms where sensory information would be integrated in a *supra-additive* manner at an early stage, while at a later stage reward can modulate the neuronal responses *sub-additively* (**model B** in Figure 1). Then, how do the two mechanisms interact with each other? To test this question, we looked into a conjunction contrast between reward effect across all sensory modalities and the *supra-additive* multisensory sensory integration. Interestingly, the conjunction revealed an area in the Fusiform Gyrus (**Figure 5D**). The coordinates of the activations in the Fusiform Gyrus found in our study overlap with areas in the Fusiform Gyrus that have been shown to be responsible for the integration of face and voice stimuli into a coherent speech perception (De Gelder & Bertelson, 2003; Rüsseler, Ye, Gerth, Szycik, & Münte, 2018) and integration of face and haptic stimuli (Kitada, Johnsrude, Kochiyama, & Lederman, 2009). In fact, it has been suggested that Fusiform Gyrus may be a convergence point across multimodal cues (Stevenson et al., 2009). In relation to reward processing, Padmala and Pessoa (2011) showed that the Fusiform Gyrus mediated the motivational cues to reduce a conflict-related processing of a target. Extending this view, Rothkirch, Schmack, Deserno, Darmohray, & Sterzer (2014) argued in their study that the Fusiform gyrus is an area where reward and attention compete for sensory processing resources. Based on our results and previous studies, we speculate the role of the Fusiform gyrus in our paradigm to act as a *hub* where sensory and reward signals converge. As fusiform gyrus did not exhibit the strongest supra-additive response to the multisensory stimuli, this might indicate that reward assists multisensory integration only when the sensory integration is weak. Another possibility is that the *supra-additive* sensory integration found in the Fusiform gyrus might occur earlier than the modulation by reward, which would be interesting to investigate using a method that has a better time resolution such as electroencephalography (EEG) or magnetoencephalography (MEG).

Moreover, we also observed areas in the premotor cortex that were modulated by both reward and multisensory integration as indexed by the conjunction contrast. Premotor cortex is an area that is critical for goal-directed behavior (Gremel & Costa, 2013) and its responses can be modulated by rewards (Peterson & Seger, 2013; Roesch & Olson, 2003). Furthermore, premotor cortex has previously been reported to be involved in multisensory integration, as it has efferent connections from multisensory areas such as the superior colliculus (Meredith, Nemitz, & Stein, 1987). The modulation of the premotor cortex revealed by the conjunction contrast may reflect that the motor preparation signals are enhanced by multimodal stimuli and this effect is further boosted by reward. Our conjunction analysis revealed both Fusiform and the premotor cortex that can be mapped to afferent and efferent communications to the brain areas that implement the final motor response. Thus, as both areas were detected using the same conjunction analysis, the Fusiform areas might process both sensory and reward information that is further signaled to the premotor areas to optimize goal-directed behavior (i.e. “how fast should I press the button?”) depending on the reward at stake.

Collectively, our behavioral and neural findings contradicted our hypothesis that reward would enhance multisensory integration according to a *supra-additive* principle. Instead, the evidence we found indicates that reward enhances multisensory integration at a late stage and *sub-additively*. Following this hypothesis, we considered possible mechanisms of the interaction between reward and multisensory integration as shown in **Figure 1**. These mechanisms either map to an *early integration scheme*, where reward is integrated as early as the sensory information is (either at the primary sensory areas or in the heteromodal areas, hence **model A and C** in Figure 1), thus this would mean that the reward effect would be *supra-additive*. In contrast, *late integration* proposed that reward information is integrated after sensory integration occurs, thus the reward effect may have a *sub-additive* pattern (**model B** in Figure 1). Lastly, the *parallel processing scheme* (**model C** in Figure 1) suggests that reward and sensory systems share similar mechanisms and can be integrated at both early and late stages (see also Koelewijn et al., 2010). Based on the evidence provided by the univariate analysis of the fMRI data, we ruled out *early integration* and *parallel processing* mechanisms, as they both predict a *supra-additive* reward modulation for multisensory stimuli. Furthermore, we observed distinct regions that exhibited strong modulations either by reward (frontal and midbrain areas) or by multisensory information (occipital and temporal regions). Thus, we showed that reward and sensory integration are two independent mechanisms, where sensory cues are integrated automatically, and then reward may enhance the integration further in a *sub-additive* manner. This indicates that reward interacts with multisensory integration at a later stage, supporting the *late integration* framework (**panel B** in Figure 1), at least in the context of the task we used in this study. Our findings are in line with early studies on the role of attention in multisensory integration, as observed in Vroomen, Bertelson, & De Gelder, (2001), where they demonstrated that cross-modal interactions such as the one observed in visual Ventriloquism effect do not depend on visual spatial attention. Furthermore, an ERP study by Santangelo, Van Der Lubbe, Olivetti Belardinelli, and Postma (2008) also revealed that although multisensory cues showed a supra-additive effect, this pattern did not extend to the cueing effect, since the cueing effect in the multisensory cues did not differ from that of unimodal cues. So can we conclude that reward, akin to some aspects of attentional processing, may not have a strong effect on multisensory integration, which largely occurs automatically?

Before the above claim about a lack of interaction between reward and multisensory processing or the existence of a late integration mechanism can be made, there are some alternative possibilities that need to be considered. In our paradigm, we marked the rewarded and not rewarded cues by presenting on the display either some amount of money (e.g. ‘20 Cent’) or none (i.e., ‘00 Cent’), respectively. However, if participants missed a cue, the same feedback display as not rewarded conditions (‘00 Cent’) was shown. Although in our experiments, cues were rarely missed (**Figure S1.A**), this setting was suboptimal as the feedback for accuracy and rewards were identical. Potentially, this might disrupt the association of the cues with the reward magnitude. Moreover, as mentioned in relation to the differences between our design and that of Bean and colleagues (Bean et al., 2021), using weak unisensory stimuli may enhance the multisensory integration and potentially also its modulation by rewards, according to the *inverse effectiveness* (but also see **Figure S1.C**). In future studies, this possibility can be investigated by presenting stimuli at or below the detection threshold for each participant. Furthermore, another consideration is the mode of reward administration. In our paradigm, we manipulated the magnitude of rewards. However, reward expectancy constitutes of both magnitude and probabilities of obtaining the reward (Schultz, 2006; Yacubian et al., 2007). For instance, future studies could investigate whether enhancing the uncertainty in the reward predictability, for instance by varying the reward probability, can have an impact on its role in multisensory integration.

To conclude, our study is the first to provide evidence for a late integration model of reward and multisensory processing. We confirmed previous findings that the information from two modalities are integrated *supra-additively*. Additionally, we showed that at this stage, reward does not influence multisensory integration. Instead, we provided evidence that reward modulation occurs at a later stage and in a *sub-additive* manner. Importantly, we found that association areas in the Fusiform gyrus show both multisensory supra-additivity and reward modulation and may hence play a role in the late integration of the two types of information.

## Acknowledgements

We thank Prof. Melanie Wilke and Dr. Roberto Goya-Maldonado for the discussions, Nina Franziska Kirchhof, Gökberk Günaydin and Veronica Wardhani for their help with the data collection and analysis. This work was supported by an ERC Starting Grant (no: 716846) to AP.

## Authors’ contributions

JEA and AP conceptualized the project designed the task. JEA conducted the experiments. JEA and AP analyzed the data. JEA and AP interpreted the results and wrote the first draft of the manuscript. All authors revised the manuscript. AP acquired funding.

## Notes

**<u>Conflict of interests:</u>** The authors declare no competing interests.

### Competing Interest Statement

The authors have declared no competing interest.

## References

Algazi, V. R., Duda, R. O., Thompson, D. M., & Avendano, C. (2001). The CIPIC HRTF database. IEEE ASSP Workshop on Applications of Signal Processing to Audio and Acoustics, (October), 99–102. https://doi.org/10.1109/aspaa.2001.969552

Bean, N. L., Stein, B. E., & Rowland, B. A. (2021). Stimulus value gates multisensory integration. European Journal of Neuroscience, 53(9), 3142–3159. https://doi.org/10.1111/ejn.15167

Beauchamp, M. S. (2005). See me, hear me, touch me: Multisensory integration in lateral occipital-temporal cortex. Current Opinion in Neurobiology, 15(2), 145–153. https://doi.org/10.1016/j.conb.2005.03.011

Bruns, P., Maiworm, M., & Röder, B. (2014). Reward expectation influences audiovisual spatial integration. Attention, Perception, and Psychophysics, 76(6), 1815–1827. https://doi.org/10.3758/s13414-014-0699-y

Calvert, G. A., Hansen, P. C., Iversen, S. D., & Brammer, M. J. (2001). Detection of audio-visual integration sites in humans by application of electrophysiological criteria to the BOLD effect. NeuroImage, 14(2), 427–438. https://doi.org/10.1006/nimg.2001.0812

Cheng, F. P. H., Saglam, A., André, S., & Pooresmaeili, A. (2020). Cross-modal integration of reward value during oculomotor planning. ENeuro, 7(1), 1–14. https://doi.org/10.1523/ENEURO.0381-19.2020

Colonius, H., & Diederich, A. (2017). Measuring multisensory integration: From reaction times to spike counts. Scientific Reports, 7(1), 1–11. https://doi.org/10.1038/s41598-017-03219-5

Cornelissen, F. W., Peters, E. M., & Palmer, J. (2002). The Eyelink Toolbox: Eye tracking with MATLAB and the Psychophysics Toolbox. Behavior Research Methods, Instruments, and Computers, 34(4), 613–617. https://doi.org/10.3758/BF03195489

De Gelder, B., & Bertelson, P. (2003). Multisensory integration, perception and ecological validity. Trends in Cognitive Sciences, 7(10), 460–467. https://doi.org/10.1016/j.tics.2003.08.014

Degerman, A., Rinne, T., Pekkola, J., Autti, T., Jääskeläinen, I. P., Sams, M., & Alho, K. (2007). Human brain activity associated with audiovisual perception and attention. NeuroImage, 34(4), 1683–1691. https://doi.org/10.1016/j.neuroimage.2006.11.019

Delong, P., & Noppeney, U. (2021). Semantic and spatial congruency mould audiovisual integration depending on perceptual awareness. Scientific Reports, 11(1), 1–14. https://doi.org/10.1038/s41598-021-90183-w

Diederich, A. (1995). Intersensory facilitation of reaction time: Evaluation of counter and diffusion coactivation models. Journal of Mathematical Psychology. https://doi.org/10.1006/jmps.1995.1020

Diederich, A., & Colonius, H. (2004). Bimodal and trimodal multisensory enhancement: Effects of stimulus onset and intensity on reaction time. Perception and Psychophysics, 66(8), 1388–1404. https://doi.org/10.3758/BF03195006

Ferenczi, E. A., Zalocusky, K. A., Liston, C., Grosenick, L., Warden, M. R., Amatya, D., … Deisseroth, K. (2016). Prefrontal cortical regulation of brainwide circuit dynamics and reward-related behavior. Science, 351(6268). https://doi.org/10.1126/science.aac9698

Ferrari, A., & Noppeney, U. (2021a). Attention controls multisensory perception via 2 distinct mechanisms at different levels of the cortical hierarchy. PLoS Biology (Vol. 19). https://doi.org/10.1371/journal.pbio.3001465

Ferrari, A., & Noppeney, U. (2021b). Attention controls multisensory perception via 2 distinct mechanisms at different levels of the cortical hierarchy. PLoS Biology (Vol. 19). https://doi.org/10.1371/journal.pbio.3001465

Gremel, C. M., & Costa, R. M. (2013). Premotor cortex is critical for goal-directed actions. Frontiers in Computational Neuroscience, 7(AUG), 1–8. https://doi.org/10.3389/fncom.2013.00110

Hoofs, V., Grahek, I., Boehler, C. N., & Krebs, R. M. (2022). Guiding spatial attention by multimodal reward cues. Attention, Perception, and Psychophysics, 84(3), 655–670. https://doi.org/10.3758/s13414-021-02422-x

Kitada, R., Johnsrude, I. S., Kochiyama, T., & Lederman, S. J. (2009). Functional specialization and convergence in the occipito-temporal cortex supporting haptic and visual identification of human faces and body parts: An fMRI study. Journal of Cognitive Neuroscience, 21(10), 2027–2045. https://doi.org/10.1162/jocn.2009.21115

Koelewijn, T., Bronkhorst, A., & Theeuwes, J. (2010). Attention and the multiple stages of multisensory integration: A review of audiovisual studies. Acta Psychologica, 134(3), 372–384. https://doi.org/10.1016/j.actpsy.2010.03.010

Lewis, J. W., & Van Essen, D. C. (2000). Corticocortical connections of visual, sensorimotor, and multimodal processing areas in the parietal lobe of the macaque monkey. Journal of Comparative Neurology, 428(1), 112–137. https://doi.org/10.1002/1096-9861(20001204)428:1<112::AID-CNE8>3.0.CO;2-9

Linden, J. F., Grunewald, A., & Andersen, R. A. (1999). Responses to auditory stimuli in macaque lateral intraparietal area II. Behavioral modulation. Journal of Neurophysiology, 82(1), 343–358. https://doi.org/10.1152/jn.1999.82.1.343

Lunghi, C., Verde, L. Lo, & Alais, D. (2017). Touch accelerates visual awareness. I-Perception, 8(1). https://doi.org/10.1177/2041669516686986

Meredith, M. A., Nemitz, J. W., & Stein, B. E. (1987). Determinants of multisensory integration in superior colliculus neurons. I. Temporal factors. The Journal of Neuroscience: The Official Journal of the Society for Neuroscience, 7(10), 3215–3229. https://doi.org/10.1523/jneurosci.07-10-03215.1987

Meredith, M. A., & Stein, B. E. (1986). Visual, auditory, and somatosensory convergence on cells in superior colliculus results in multisensory integration. Journal of Neurophysiology, 56(3), 640–662. https://doi.org/10.1152/jn.1986.56.3.640

Nagy, A., Eördegh, G., Paróczy, Z., Márkus, Z., & Benedek, G. (2006). Multisensory integration in the basal ganglia. European Journal of Neuroscience, 24(3), 917–924. https://doi.org/10.1111/j.1460-9568.2006.04942.x

Nagy, A., Paróczy, Z., Norita, M., & Benedek, G. (2005). Multisensory responses and receptive field properties of neurons in the substantia nigra and in the caudate nucleus. European Journal of Neuroscience, 22(2), 419–424. https://doi.org/10.1111/j.1460-9568.2005.04211.x

Noppeney, U. (2012). Characterization of Multisensory Integration with fMRI: Experimental Design, Statistical Analysis, and Interpretation. CRC Press/Taylor & Francis, Boca Raton (FL).

Otto, T. U., Dassy, B., & Mamassian, P. (2013). Principles of multisensory behavior. Journal of Neuroscience, 33(17), 7463–7474. https://doi.org/10.1523/JNEUROSCI.4678-12.2013

Padmala, S., & Pessoa, L. (2011). Reward reduces conflict by enhancing attentional control and biasing visual cortical processing. Journal of Cognitive Neuroscience, 23(11), 3419–3432. https://doi.org/10.1162/jocn_a_00011

Pessoa, L., & Engelmann, J. B. (2010). Embedding reward signals into perception and cognition. Frontiers in Neuroscience, 4(SEP). https://doi.org/10.3389/fnins.2010.00017

Peterson, E. J., & Seger, C. A. (2013). Many hats: Intratrial and reward level-dependent BOLD activity in the striatum and premotor cortex. Journal of Neurophysiology, 110(7), 1689–1702. https://doi.org/10.1152/jn.00164.2012

Reig, R., & Silberberg, G. (2014). Multisensory Integration in the Mouse Striatum. Neuron, 83(5), 1200–1212. https://doi.org/10.1016/j.neuron.2014.07.033

Renier, L. A., Anurova, I., De Volder, A. G., Carlson, S., VanMeter, J., & Rauschecker, J. P. (2009). Multisensory integration of sounds and vibrotactile stimuli in processing streams for “what” and “where.” Journal of Neuroscience, 29(35), 10950–10960. https://doi.org/10.1523/JNEUROSCI.0910-09.2009

Roesch, M. R., & Olson, C. R. (2003). Impact of expected reward on neuronal activity in prefrontal cortex, frontal and supplementary eye fields and premotor cortex. Journal of Neurophysiology, 90(3), 1766–1789. https://doi.org/10.1152/jn.00019.2003

Romei, V., Murray, M. M., Merabet, L. B., & Thut, G. (2007). Occipital transcranial magnetic stimulation has opposing effects on visual and auditory stimulus detection: Implications for multisensory interactions. Journal of Neuroscience, 27(43), 11465–11472. https://doi.org/10.1523/JNEUROSCI.2827-07.2007

Rothkirch, M., Schmack, K., Deserno, L., Darmohray, D., & Sterzer, P. (2014). Attentional modulation of reward processing in the human brain. Human Brain Mapping, 35(7), 3036–3051. https://doi.org/10.1002/hbm.22383

Rüsseler, J., Ye, Z., Gerth, I., Szycik, G. R., & Münte, T. F. (2018). Audio-visual speech perception in adult readers with dyslexia: an fMRI study. Brain Imaging and Behavior, 12(2), 357–368. https://doi.org/10.1007/s11682-017-9694-y

Santangelo, V., Van Der Lubbe, R. H. J., Olivetti Belardinelli, M., & Postma, A. (2008). Multisensory integration affects ERP components elicited by exogenous cues. Experimental Brain Research, 185(2), 269–277. https://doi.org/10.1007/s00221-007-1151-5

Schroeder, C. E., & Foxe, J.. (2002). Schroeder02, 14, 1–12.

Schultz, W. (2000) Multiple reward signals in the brain. Nat Rev 1:199–207.

Schultz, W. (2006). Behavioral theories and the neurophysiology of reward. Annual Review of Psychology, 57, 87–115. https://doi.org/10.1146/annurev.psych.56.091103.070229

Senkowski, D., Talsma, D., Herrmann, C. S., & Woldorff, M. G. (2005). Multisensory processing and oscillatory gamma responses: effects of spatial selective attention. Experimental Brain Research, 166(3–4), 411–426. https://doi.org/10.1007/s00221-005-2381-z

Soto-Faraco, S., Kingstone, A., Calvert, G., Spence, C., & Stein, B. E. (2004). Multisensory Integration of Dynamic Information. The Handbook of Multisensory Processes, (1981), 1–63.

Stein, B. E., Meredith, M. A., & Wallace, M. T. (1993). The visually responsive neuron and beyond: Multisensory integration in cat and monkey. Progress in Brain Research, 95(C), 79–90. https://doi.org/10.1016/S0079-6123(08)60359-3

Stein, B. E., & Stanford, T. R. (2008). Multisensory integration: Current issues from the perspective of the single neuron. Nature Reviews Neuroscience, 9(4), 255–266. https://doi.org/10.1038/nrn2331

Stein, B. E., Stanford, T. R., Ramachandran, R., Jr, T. J. P., & Rowland, B. A. (2009). Challenges in quantifying multisensory integration. Experimental Brain Research, 198(2–3), 113–126. https://doi.org/10.1007/s00221-009-1880-8.Challenges

Stevenson, R. A., Kim, S., & James, T. W. (2009). An additive-factors design to disambiguate neuronal and areal convergence: Measuring multisensory interactions between audio, visual, and haptic sensory streams using fMRI. Experimental Brain Research, 198(2–3), 183–194. https://doi.org/10.1007/s00221-009-1783-8

Talsma, D., & Woldorff, M. G. (2005). <Talsma_JoCN_2005.pdf>, 1098–1114.

Van der Stoep, N., Van der Stigchel, S., Van Engelen, R. C., Biesbroek, J. M., & Nijboer, T. C. W. (2019). Impairments in multisensory integration after stroke. Journal of Cognitive Neuroscience, 31(6), 885–899.

Vroomen, J., Bertelson, P., & De Gelder, B. (2001). The ventriloquist effect does not depend on the direction of automatic visual attention. Perception and Psychophysics, 63(4), 651–659. https://doi.org/10.3758/BF03194427

Wallace, M. T., Meredith, M. A., & Stein, B. E. (1993). Converging influences from visual, auditory, and somatosensory cortices onto output neurons of the superior colliculus. Journal of Neurophysiology, 69(6), 1797–1809. https://doi.org/10.1152/jn.1993.69.6.1797

Wallace, Mark T., & Stein, B. E. (1997). Development of multisensory neurons and multisensory integration in cat superior colliculus. Journal of Neuroscience, 17(7), 2429–2444. https://doi.org/10.1523/jneurosci.17-07-02429.1997

Werner, S., & Noppeney, U. (2010). Distinct functional contributions of primary sensory and association areas to audiovisual integration in object categorization. Journal of Neuroscience, 30(7), 2662–2675. https://doi.org/10.1523/JNEUROSCI.5091-09.2010

Werner, S., & Noppeney, U. (2011). The contributions of transient and sustained response codes to audiovisual integration. Cerebral Cortex, 21(4), 920–931. https://doi.org/10.1093/cercor/bhq161

Yacubian, J., Sommer, T., Schroeder, K., Gläscher, J., Braus, D. F., & Büchel, C. (2007). Subregions of the ventral striatum show preferential coding of reward magnitude and probability. NeuroImage, 38(3), 557–563. https://doi.org/10.1016/j.neuroimage.2007.08.007

