## Supplemental Information for "Modulation of perception by visual, auditory, and audiovisual reward predicting cues"

### Supplementary Information

#### Caudate BOLD response to *supra-additive* sensory integration and reward predicting cues

To investigate further whether the areas of the Caudate responding to the *supra-additive* sensory integration overlap with the areas that were modulated by reward, we extracted and overlaid the two functional ROIs (see **Figure S1**). The areas in the Caudate showing the reward modulation (xyz = [9 11 -1]) and *supra-additive* sensory integration (xyz = [15 14 20]) did not overlap, suggesting that areas in the Caudate had different functionality. In fact, reward modulation was observed in ventral caudate in line with previous observations (Nakamura et al., 2012), whereas integration of multimodal sensory inputs was observed in dorsal Caudate (Haber, 2011).

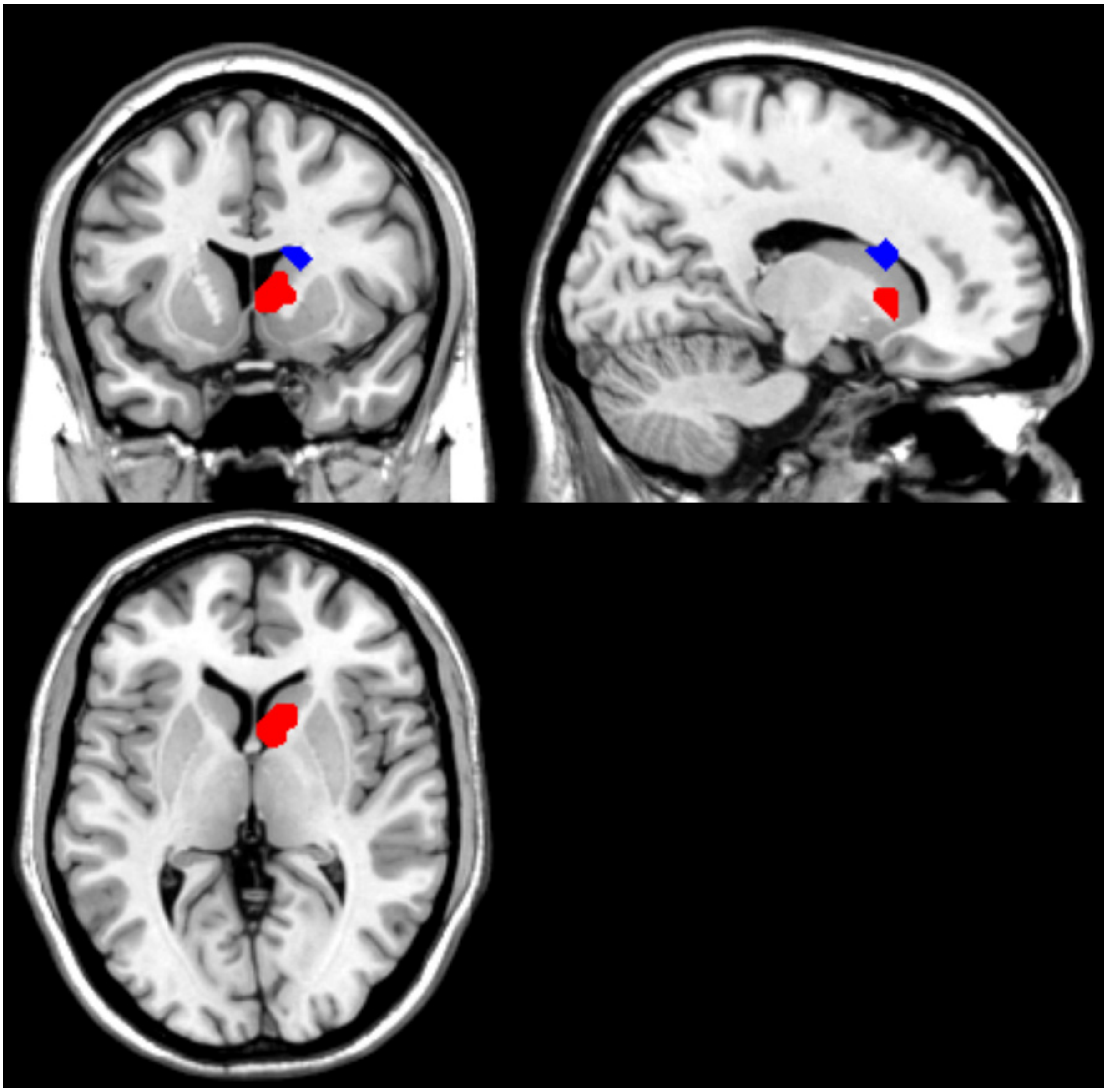

**Figure S1.** Overlays of the Caudate activities in reward modulation across sensory modalities (in red), where the peak is at xyz = [9 11 -1] and *supra-additive* sensory integration (in blue), where the peak is at xyz = [15 14 20]. ROIs were obtained from images thresholded at uncorrected $p < 0.001$ with extent threshold 10. Cursor is at xyz = [15 11 4].

#### Reward effects on the strongest areas in the supra-additive sensory integration

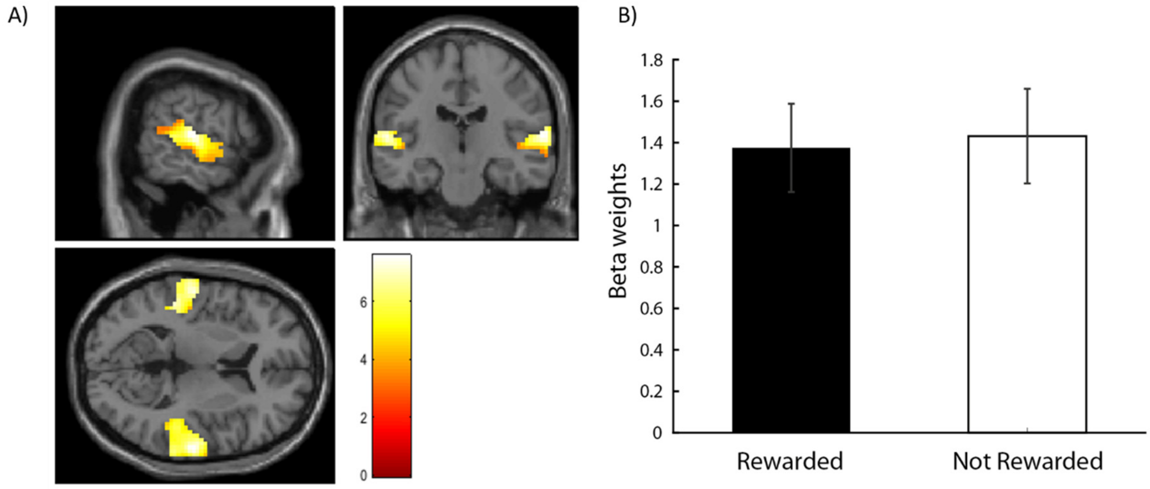


**Figure S2.** A) Regions of interest (ROI) of superior temporal areas extracted from the supra-additive sensory integration contrast (thresholded at uncorrected $p < 0.001$, k = 10) and masked with an anatomical superior temporal areas from the AAL atlas. Cursor is located at the global maximum xyz=[63 -19 8]. B) Functional ROI analysis examining reward effects on the beta weights of the superior temporal ROI.

We examined reward effects using paired sample *t*-test of the functional region of interest (fROI) analysis in the strongest areas showing *supra-additive* sensory integration in the superior temporal (peak at xyz=[63 -19 8], **Figure S2A**). However, reward did not modulate the activities in the superior temporal areas as demonstrated by the paired sample *t*-test ($p$ = 0.47, Cohen's d = 0.15, **Figure S2B**).

#### Identification of brain areas that showed a reverse reward modulation

Since reward can either enhance or suppress neural responses, we examined the reverse reward contrasts, i.e. High Reward < Low Reward, for all comparisons that are reported in the main text. This analysis predominantly revealed areas in the frontal and fronto-occipital areas as shown in **Table S1**.

**Table S1**. Whole-brain activations of univariate results thresholded at uncorrected $p < .001$ and k = 10 for the inversed reward-effect (Rewarded<not Rewarded). Significance ($p$) are reported for cluster-level.

| Cluster size | MNI coordinates (in mm) | | | T | *p* | Side | Region |
|---|---|---|---|---|---|---|---|
| | x | y | z | | | | |
| Inversed main effect of reward (AVnot rew+Anot rew+Vnot rew > AVrew+Arew+Vrew) | | | | | | | |
| 107 | -30 | 35 | 44 | 4.48 | 0.028 | L | Middle frontal |
| 215 | -45 | -58 | 23 | 4.16 | 0.003 | L | Angular |
| 77 | -3 | -58 | 23 | 3.84 | 0.056 | L | Precuneus |
| 26 | -63 | -10 | -22 | 3.63 | 0.248 | L | Middle temporal |
| 13 | 27 | 35 | 41 | 3.58 | 0.416 | R | Middle frontal |

| | | | | | | | |
|---|---|---|---|---|---|---|---|
| Inversed reward effect in audiovisual (AVnot rew > AVrew) | | | | | | | |
| 194 | -27 | 35 | 41 | 4.33 | 0.005 | L | Middle frontal |
| 50 | -48 | -55 | 29 | 3.61 | 0.116 | L | Angular |
| 16 | 27 | 32 | 44 | 3.51 | 0.365 | R | Middle frontal |
| 14 | -60 | -13 | -22 | 3.45 | 0.397 | L | Middle temporal |
| Inversed reward effect in visual (Vnot rew > Vrew) | | | | | | | |
| 28 | -3 | -55 | 23 | 3.46 | 0.231 | L | Precuneus |
| 17 | -42 | -61 | 26 | 3.43 | 0.35 | L | Angular |
| Inversed reward effect in auditory (Anot rew > Arew) | | | | | | | |
| *No voxels survived* | | | | | | | |

#### References


Haber SN (2011) 11 neuroanatomy of reward: A view from the ventral striatum. Neurobiol Sensat Reward:235.

Nakamura K, Santos GS, Matsuzaki R, Nakahara H (2012) Differential reward coding in the subdivisions of the primate caudate during an oculomotor task. J Neurosci 32:15963–15982.

Stein BE, Stanford TR (2008) Multisensory integration: Current issues from the perspective of the single neuron. Nat Rev Neurosci 9:255–266.
